# Stability of Aβ-fibril fragments in the presence of fatty acids

**DOI:** 10.1101/620518

**Authors:** Wenhui Xi, Elliott K. Vanderford, Qinxin Liao, Ulrich H. E. Hansmann

**Author notes:** **Corresponding Author** Ulrich H. E. Hansmann.

## Abstract

We consider the effect of lauric acid on the stability of various fibril-like assemblies of Aβ peptides. For this purpose, we have performed molecular dynamics simulations of these assemblies either in complex with lauric acid or without presence of the ligand. While we do not observe a stabilizing effect on Aβ_40_-fibrils we find that addition of lauric acid strengthen the stability of fibrils built from the more toxic triple-stranded S-shaped Aβ_42_-peptides. Or results may help to understand how specifics of the brain-environment modulate amyloid formation and propagation.

## Introduction

An increasing list of diseases is characterized by the presence of amyloid fibrils visible under ultraviolet light after staining with Congo Red.^1^ Probably the best-known example is Alzheimer’s disease, a neurodegenerative illness mostly striking the elderly, where in a postmortem patient brains are characterized by amyloid deposits rich in Aβ peptides.^2^ Neither the disease mechanism nor the exact agent is known, but considerable evidence points to soluble oligomers instead of the insoluble fibrils as the more toxic species.^3-5^

Both oligomers and fibrils are *in vitro* characterized by a rich structural polymorphism. However, the same diversity of structures is not seen in fibrils derived from patient brains^6^, which instead have a unique, patient-specific, form. We have shown in earlier work that this lack of polymorphism cannot be explained by a higher thermodynamic stability of the patient-derived fibril structure^7^. A more likely scenario appears to be the spread of “infectious” strains, i.e. certain amyloid structures may prion-like self-replicate and propagate^8-10^, a conjecture that is supported by experiments with mice, and *in vitro* for fibrils formed by the Osaka mutant.^11-13^

While this scenario would explain the observed lack in fibril diversity seen in patient brains, it opens up the question of why the same process is not observed *in vitro*. A possible explanation may be the brain-specific environment. For instance, lipid and fatty acid interfaces can enhance oligomer formation preferentially over fibrils^14-16^. Examples are glycolipid-rich Golgi membranes that induce formation of oligomer but not of fibrils^17^; and lipid rafts isolated from brain tissue result only in the assembly of tetramers^16^. Fatty acids also have significant effects on Alzheimer patient brains^18^, and even nanomolar concentrations of Aβ can form pathological aggregates *in vivo* that are not observed *in vitro*^*19,20*^. Hence, one could hypothesize that lipid and fatty acid interfaces encourage the formation of self-replicating and self-propagating Aβ-assemblies, which finally lead to the unique fibril structures observed in patient brains. This hypothesis is supported by the observation of prion-like self-propagating Large Fatty Acid-derived Oligomers (LFAOs), generated in the presence of lauric acid and stable after removal of the fatty acids, that in mice are causing Cerebral Amyloid Angiopathy (a common co-pathology in Alzheimer’s disease patients).^21^

The above described experiments have motivated us to study in this paper how the presence of lauric acid alters the stability of Aβ-fibrils. The first model that we consider is the sole patient-derived fibril model, which is built from U-shaped Aβ_40_-peptides and has been deposited in the Protein Data Bank under PDB-Id 2M4J. However, the more cytotoxic Aβ_42_-peptides can assume three-stranded S-shaped folds, enabling them to aggregate into assemblies that cannot be formed by the U-shaped Aβ_40_. Specifically, we have argued in previous work that the three-stranded S-shaped Aβ_42_-peptides can assemble into ring-like oligomers and fibrils^22^, and one of these assemblies, a double-layered hexamer, was found to be consistent with LFAO measurements.^23^ For this reason, we study here also how the stability of such Aβ_42_-assemblies is affected by the presence of lauric acid. Considering these diverse systems may help us to understand how lipid and fatty acid interfaces in the brain influence amyloid formation and propagation.

## Methods

### Model construction

We probe the effect of lauric acid on the stability of Aß_40_ fibrils by considering the single patientderived fibril model deposited in the Protein data Bank (PDB-ID:2M4J).^6^ This ss-NMR resolved structure has a three-fold symmetry, and is built from U-shaped Aß_40_ chains with an in-register parallel β-sheet made of two strands, residues 8-20 and 29-34. We have studied this model in earlier work,^7^ and found the six-layer fragment to be only marginal stable, with considerable differences in stability between the 20 NMR models deposited in the PDB under the PDB-ID 2M4J. For this reason, we chose in the present work a four-layer fragment which we know would not be stable by itself. Taking the ensemble of NMR structures as a rough approximation of the equilibrium ensemble of fibril configurations, each allowing or preventing binding in slightly different ways, we have predicted binding sites for lauric acid for all 20 models by AutoDockVina.^24^ For this purpose, we take the lauric acid structure from the PubMed Compound database.^25^ From the set of predicted binding sites, we exclude the ones located on the extension surfaces of fibrils as we are not interested in the influence of the ligand on fibril formation or elongation. A visual analysis of the remaining complexes allowed us identify four possible binding sites: two involving the N-terminal residues 1-8, a cave in the turn-loop region of residues 21-30, and the C-terminal residues 31-36 and 39-40. For each binding site we chose the NMR model with the lowest energy (and presumably highest stability), i.e., the one with lowest model index, see Fig. 1(A). Hence, starting point for our simulations are the NMR models 1,2,8, and 20 in complex with lauric acid. The four NMR-models are shown in Fig. 1(B) – 1(E), while the lauric acid is displayed in subfigure 1(A).

**Figure 1.**
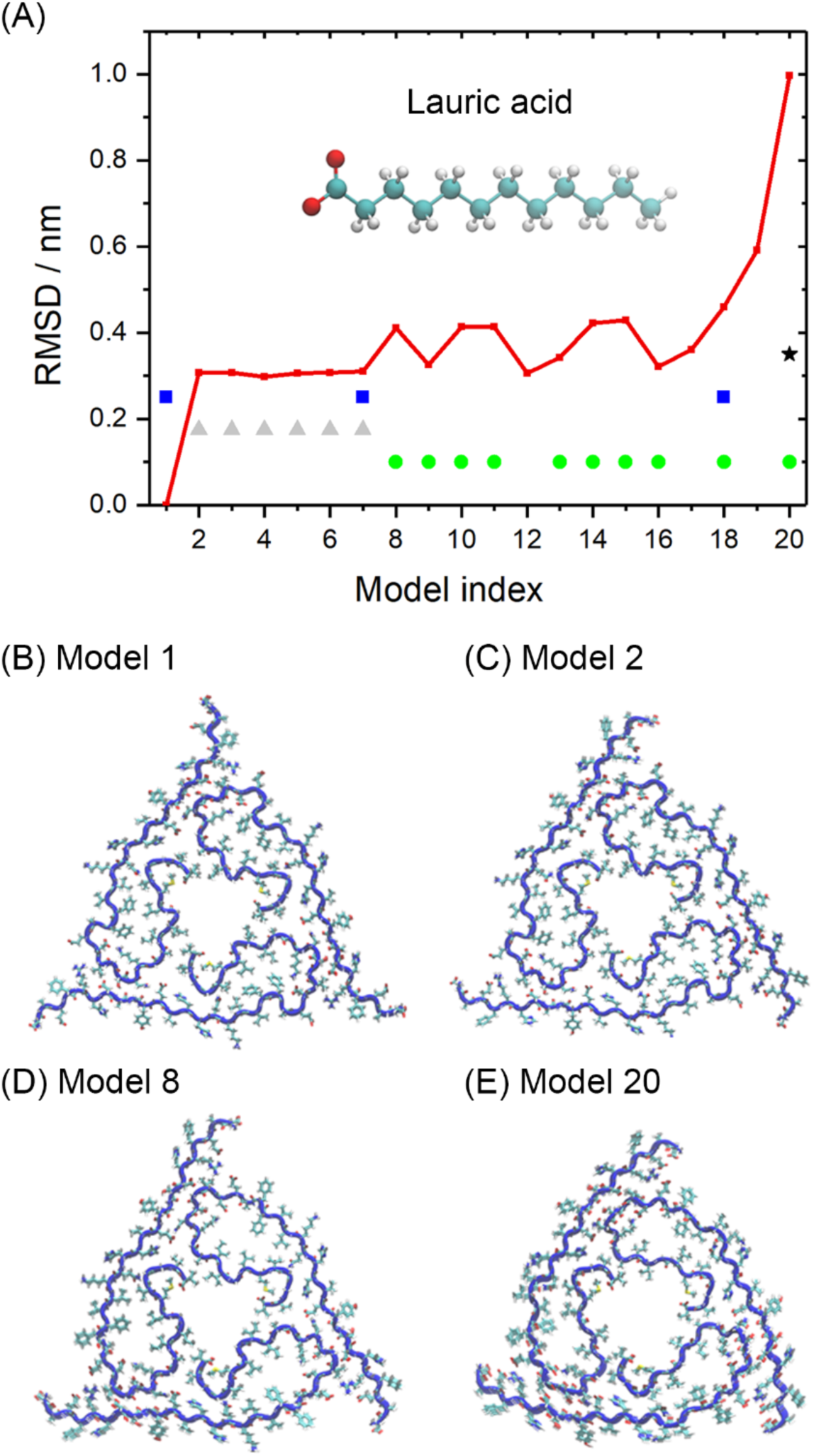
The root-mean-square deviation (RMSD) in relation to the first model for all 20 models of the patient-derived Aβ_40_ fibril, deposited in the Protein data Bank under identifier 2M4J, is shown in sub-figure (A). Four distinct binding sites were found. The grey triangle marks binding to N-terminal residues with odd index (5,7,9), the blue square marks to N-terminal residues with even indices (6,8,10); a green circle marks binding to a cave in the turn-loop region of residues 21-30; and a black star marks binding to C-terminal residues. Sub-figure (A) shows also the initial configuration of the lauric acid molecules. For our investigation we chose for each binding site the model which has the lowest RMSD to the first model. The initial configurations of these four models are shown in sub-figures (B-E), with model 1 and model 2 chosen to probe the two N-terminal binding sites, model 8 to probe binding to the turn-loop cave, and model 20 for the binding to C-terminal residues.

The interaction of lauric acid with the three-stranded Aß_42_-assemblies is studied first for two-fold models of Aß_11_-_42_ with varying the number of layers, and with the binding pattern proposed by us in previous work^26^. We later consider two ring-like dodecamers, a three-layered tetramer and a two-layered hexamer which were proposed in earlier work^22^ as models for the LFAOs extensively studied by the Rangachari lab^21,27,28^. All three types of assemblies are displayed in Fig. 2 (A-C). We use again AutoDockVina^24^ to predict possible binding sites, and have identified three groups. The first binding site is between the packing surface of two folds where the lauric acid molecule interacts with residue K16 on one side and residue D22 on the other side. The second site is similar to the first, but the lauric acid interacts here with residues K16 and V18. The third site is also on the packing surface but with the lauric acid now interacting with residues 10-13 on one side and 27-30 (or 42) on the other side.

**Figure 2.**
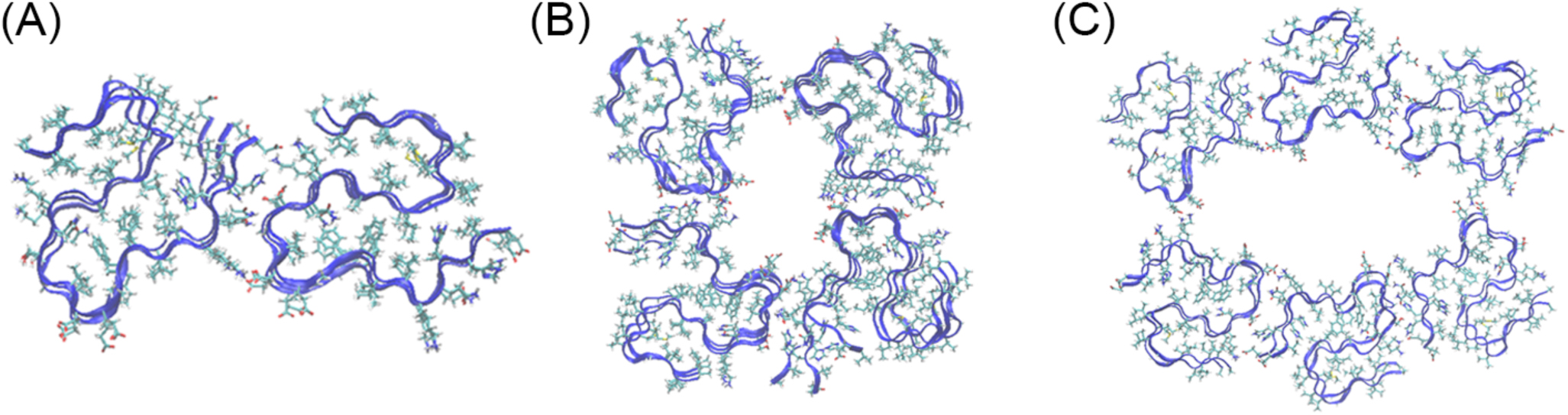
The initial conformation of the three-layer two-fold Aβ_42_ fibril model is shown in sub-figure (A), while the three-layer four-fold ring-like model is displayed in (B), and the two-layer six-fold ring-like model in (C).

### Molecular dynamics simulations

Our simulations utilize the GROMACS 5.1.5^29^ tool suit, with the Aß-peptides described by the CHARMM36 force field (version: jul-2017),^30^ and the solvent modeled by TIP3P water.^31^ The force field parameters of lauric acid are generated by the online tools *cgenff*.^32^ The temperature of 310K and a pressure of 1 bar are controlled by the v-rescale thermostat^33^ and Parrinello-Rahman barostat.^34^ Constraining bonds with the LINCE algorithm^35^ or the SETTLE algorithm^36^ allows us to use a time step of 2 fs for integration. The fibril-ligand complex and water are placed in a box with minimum distance of 0.8nm to the peptides. Use of periodic boundary conditions requires calculation of electrostatic interactions by the particle mesh Ewald(PME) method^37^. The cutoff for van der Waals (vdW) and electrostatic interactions is set to 1.2 nm, as suggested by the developers for CHARMM force field version 36. The solute is neutralized by Na+ and Cl- ions. Our start configurations are prepared by minimizing the complex-solvent system and relaxed during 5ns of molecular dynamics in an NVT ensemble, followed by 5ns in an NPT ensemble. The time evolution of each system is followed in three independent trajectories over 50 ns, starting from different initial conditions and velocity distributions. For our analysis we use the tool set provided by GROMACS, with snapshots of configurations visualized by VMD^38^.

## Results

The purpose of the present paper is to investigate whether ligands such as lauric acid, characteristic for the brain environment, may raise the stability of certain Aβ amyloids structures that could become seeds for rapid propagation of fibrils in patient brains. For this purpose, we consider first various complexes of lauric acid with the sole patient-derived fibril structure deposited in the Protein Data Bank, which is built from U-shaped Aß_40_ chains; and compare the stability of these complexes with that of fibril fragments not bound with lauric acid.

In our previous work^7^ we have studied six-layer fragments of fibril model 1 and model 20 (the one with largest RMSD to model 1) and found only marginal stability for model 1, and slightly higher stability for model 20. In our new simulations we look now into four of the 20 NMR models (the ones with model index 1,2,8 and 20), but consider only four-layer fibril fragments. The reason for the smaller number of layers is that these fragments are expected to be less stable, and therefore any stabilizing effect of added lauric acid should be easier to detect. Each model is set-up as described in the method section and followed over 50 ns in three independent runs at T=310 K and 1 bar.

As expected, the four-layer fibril fragments alone are not stable and decay along the trajectories. This can be seen from the time evolution of the root-mean-square-deviation (RMSD) in Fig. S1. Here and in the following figures we show for a given model always only values for the trajectory that leads to the largest final RMSD values (i.e., the worst case). We show not only the RMSD values for the complete fibril, but also RMSD values for the individual fold that deviates most from its initial configuration. The latter quantity allows us to evaluate the stability of single folds. The high RMSD values for the whole fibril are caused by the loss of packing surface between layers leading to dissolution of the fibril. However, the inter-chain hydrogen bonding within layers is preserved, and while the individual units become flexible they keep their secondary structure. Representative final conformations of the control runs for the four considered models (A-D), simulated without lauric acid and the corresponding time series of RMSD values are shown as supplemental Fig. S1.

When considering the stability of complexes of the models with lauric acid we look first into the case where the binding site is in the N-terminal region. Such binding sites were found for about half of the 20 models, and appear in two variants, shown in Fig. 3 (E-H). In one case, seen for model 1 in Fig. 3 (E), the ligand interacts from “outside” with even-index residue H6, while in the second case, seen in Fig. 3 (F) for model 2, the ligands bind from “inside” with odd-index residues R5 and D7. Noticing that in both cases, lauric acid interact with E3 since its side chain direction differs in model 1 and model 2. Both binding patterns do not increase the stability of the fibril fragment and the ligand quickly un-binds from the N-terminus (see Fig. 3 (E) and (F)), leading to final RMSD values that are comparable to the control, see Fig. 3 (A) and (B). Similar unstable are complexes of lauric acid with the C-terminals, as found in model 20 and shown in Fig. 3 (D, H), where the fibrils decay faster than in the control runs and binding of lauric acid even seems to decrease stability. On the other hand, when lauric acid binds to the turn-loop cave as in model 8 (see Fig. 3 (G)), the ligand stays tightly bound. While the fragment is still dissolving, with the final RMSD values in Fig. 3(C) comparable to the control, the fluctuation in individual folds decrease from 1.0nm to about 0.8nm in the complex. Hence, binding of lauric acid to the turn-loop cave stabilizes the conformation of individual units and folds, but does not alter the cohesion between the folds.

**Figure 3.**
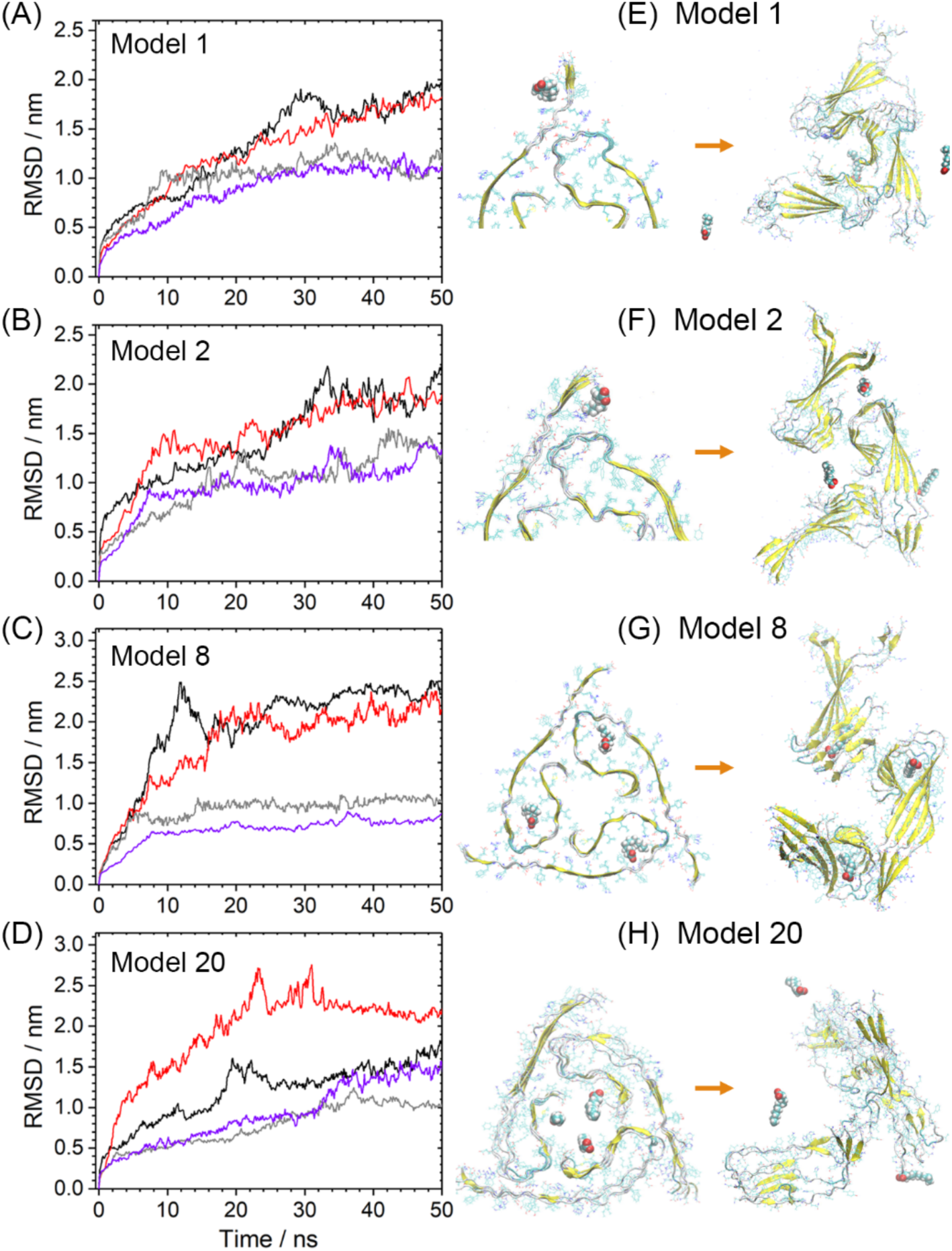
Time evolution of the four models in complex with lauric acid. In sub-figure (A) we show the root-mean-square-deviation (RMSD) of model 1 where the ligand binds to the outside of the Nterminal; in (B) that of model 2, where the ligand interacts with odd-numbered residues on the N-terminus; in (C) that of model 8 where the binding site is the turn-loop cave; and in (D) for model 20 where lauric acid binds to C-terminal residues. Results from the control run are drawn in black (whole fragment) and grey (single fold), and for the complex in red (whole fragment) and purple (single fold). The initial and final configurations for the four complexes are shown in sub-figures (E)-(H).

The above data indicate that binding of lauric acid to the patient-derived fibril structure 2M4J does not raise significantly the stability of this fibril. Cohesion between layers is not enhanced, and the sole effect appears to be a small strengthening of the fold geometry when lauric acid binds to a cave in the turn-loop region of residues 21-30. However, while 2M4J is the only patient-derived fibril structure deposited in the Protein Data Bank, it is not clear how significant this structure is for the pathogeny of Alzheimer’s disease. This is because it is believed that the more toxic assemblies are built from Aβ_42_ chains instead of the Aβ_40_ chains forming the 2M4J fibril.^39^ Unlike Aβ_40_ chains, Aβ_42_ peptides can assemble not only into fibrils built out of U-shaped chains, but also can form triple-stranded S-shaped structures.^40^ We have demonstrated in earlier work that this allows Aβ_42_ peptides to assemble into oligomer and fibril structures not accessible for Aβ_40_; and we have speculated that the enlarged structural space is related to its higher toxicity.^26^ Especially, we could construct a series of ring-like N-fold fibrils models^22^ built from S-shaped Aβ_42_ chains that allows us to propose a model for the Large-Fatty-Acid-derived - Oligomers (LFAOs) which is consistent with the low-resolution measurements of the Rangachari Lab.^23,28^ As the experiments by the Rangachari Lab have shown that fatty acids such as lauric acid can catalyze the formation of specific Aβ_42_ assemblies,^14^,^28^ we wanted to see how lauric acid affects the stability of our constructs, proposed by us as models for the disease-causing toxic oligomers.

Our models rely on a specific binding pattern between two S-shaped Aβ_42_ chains that is shown in Fig. 4 (A), where we present a segment of two chains taken from the twelve-fold model, the most stable ring-like structure found in our previous study.^22^ Our guiding hypothesis is that lauric acid catalyzes and stabilizes formation of such segments that when coagulating naturally form ring-like structures. In previous work we had seen that such two-fold assemblies are stable by themselves if made of six layers.^26^ Hence as a first step we investigated the number of layers where the stability of the two-fold assemblies is on a cusp: fewer layers will decay quickly while larger numbers of layers are stable over the investigated time scales. Any effect of lauric acid on the stability of the two-fold assemblies should be seen most clearly for this critical layer number. In order to determine this critical layer number, we therefore simulated two-fold assemblies with the binding pattern of Fig. 4 (A), built from either one, two, three, four or five layers. The RMSD to the start configuration as function of time is shown in Fig. 5 (A). This plot indicates that the critical layer size is three. If there are four or more layers, the fibrils are stable over the simulation time scale (Fig 5 (E)), while they decay quickly for less than three layers (Fig. 5 (C)) as the hydrogen bonds dissolve and the fold-assembly becomes twisted. Even in the three-layer system b the RMSD becomes larger than 1.0 nm after 50ns, with the high value resulting both from a decay of the S-shape structure of the chains in each fold, and from the relate movement between two-fold, as shown in Fig. 5 (B) and 5 (D).

**Figure 4.**
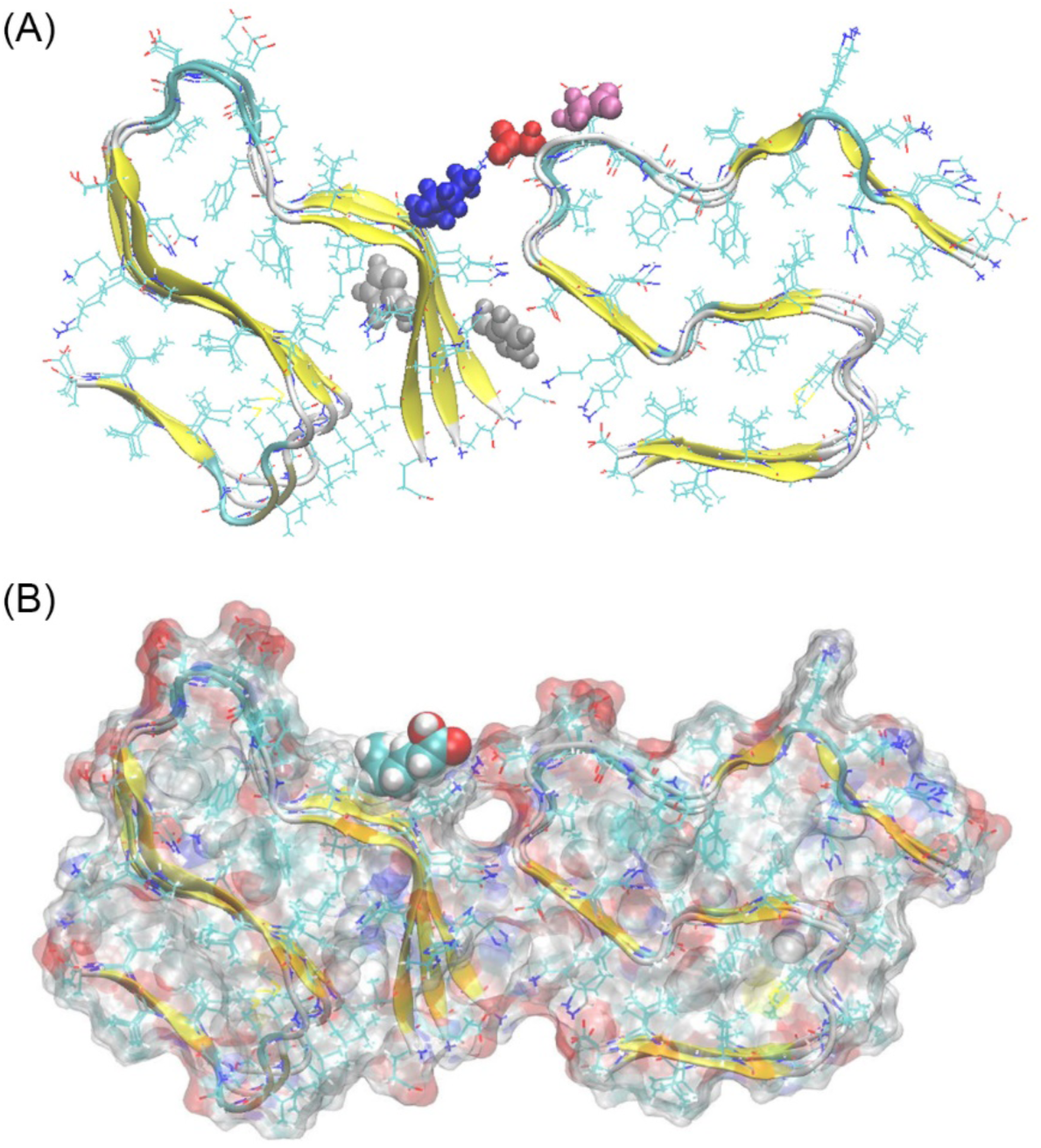
(A) Binding pattern between Aβ_42_ chains in our proposed N-fold ring-like models. Here we mark residue K16 in blue, E22 in mauve, D23 in red, and the histidines H13 and H14 in grey. The binding site of lauric acid on the three-layer model is shown in (B).

**Figure 5.**
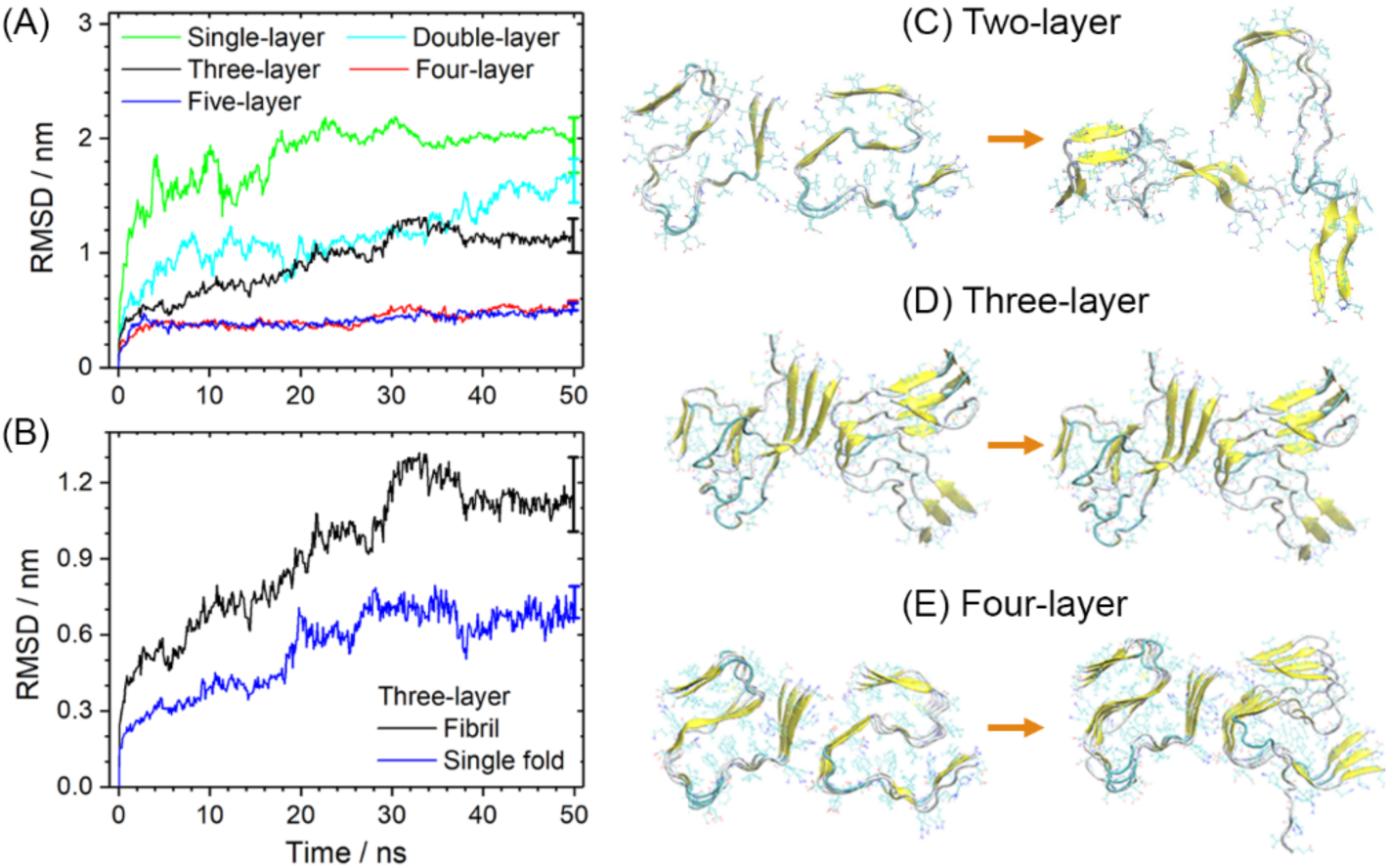
(A) The RMSD to the start configuration as function of time for our two-fold Aβ_42_ models with one to five layers. For the three-layer model, which is at the cusp of stability, we show in (B) separately the RMSD for the full system and for single folds. Shown are in both sub-figures (A) and (B) RMSD averages over the three trajectories. Initial and final configurations for three systems are shown in sub-figure (C) to (E): the unstable two-layer system, three-layer system (which is at the cusp of stability) and the stable four-layer system.

As we expect that the effect of lauric acid on the stability of the two-fold constructs is best visible for the critical layer number, we have compared in a second step the stability of three-layered assemblies bound with lauric acid with the control group of the assemblies containing no lauric acid. Each system is again followed over four independent trajectories. The three-layered two-fold assembly decays in all four trajectories for the control (i.e. in the absence, of lauric acid), see Fig. 6(A) where we show again the RMSD as function of time. A similar picture is seen in Fig. 6(B) where we show the time evolution of RMSD in four runs of a system where a single lauric acid molecule is added. While in one of the four runs the stability is remarkably higher, it is not clear whether this is a statistical fluke or indicates indeed a raised stability. However, when adding lauric acid in a ratio of 1:1, i.e., six molecules for the six Aβ_42_ peptides, the resulting complexes are clearly much more stable than the control. This can be seen for all four runs in Fig. 6(C), where the RMSD values are much lower than in the control. Note that a ratio of 1:1 was used in the experiments of the Rangachari group and found to be necessary for catalyzing the assembly of LFAO oligomers. The stabilizing effect is best seen in Fig. 6(D) where we show for both control and the complex RMSD values that are averages over four trajectories. Clearly, lauric acid not only increases the overall stability but also that of individual folds. Note that the lauric acid molecules are not tightly bound to their original binding site. Instead they can move and while most of the time staying close to the peptide surface, they can sometimes even escape into the solvent, see also Fig. 6(C) and 6(D). This behavior may also explain the concentration effect: with a ratio of 1:1 the probability of returning to their original binding sites is increased and the stabilizing effect is higher.

**Figure 6.**
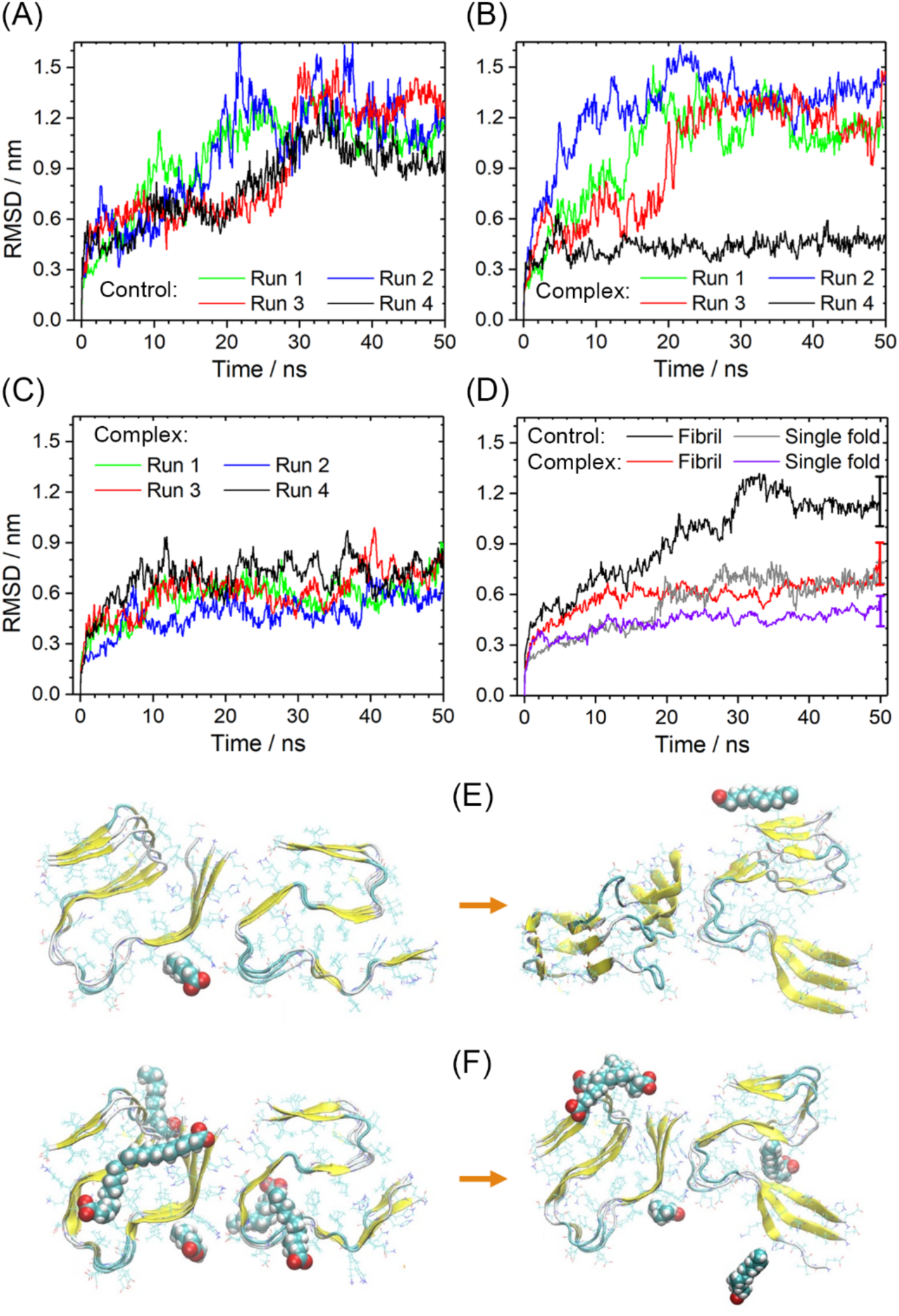
RMSD as function of time for the control model (A), the fibril binding with a single lauric acid molecule (B), and the fibril binding with six lauric acid molecules (C). For each system, data from four independent trajectories are shown. For the most stable case, the fibril binding to six molecules, we compare in (D) the average RMSD values for this complex with that of the control, considering both RMSD values for the whole system and single folds. We further show the initial and final configuration of the complex with a single lauric acid (E) and of the complex with the six-lauric acid molecules (F).

What is the effect of lauric acid molecules on the less stable single-layer and double-layer assemblies? In order to study this question, we have repeated the simulations for single-layer and double-layer two-fold assemblies bound with six lauric acid molecules. Notice that the ratio of lauric acid in single-layer (1:3) and double-layer (2:3) are higher than in the case of three layers where it is (1:1). The time evolution of the RMSD in these systems is again followed in four trajectories and compared to that of the respective control systems lacking lauric acid. The resulting plots are shown in supplemental Fig. S2, and indicate at best a marginal stabilizing effect of the lauric acid on the fibril fragments. This is expected as the flexibility of the oligomers is large: the single-layer model has no stabilizing inter-chain hydrogen bonds, and the double layer is stabilized by only by a single set of about 16 hydrogen bonds (while starting with the three-layer systems the inner chains are stabilized by two sets of hydrogen bonds, one set on each side). The resulting much smaller number of inter-chain hydrogen bonds in the single-layer and double-layer models is difficult to overcome by any potentially stabilizing effect of lauric acid binding. In the case of the single-layer model we observe not even a stabilization of the chain geometry. This may indicate that lauric acid is encouraging and supporting multi-layer assemblies but not enhancing the formation of multi-fold assemblies.

Our above results suggest that while lauric acid is not stabilizing the Aβ_40_-fibril, it has a profound and concentration-dependent effect on the stability of amyloids made from S-shaped Aβ_42_-chains. Our assumption is that the strengthening of the fibril geometry allows lauric acid to catalyze the fatty acid oligomers studied by the Rangachari Lab^28^ and named LFAOs. At low Aβ_42_ concentration these LFAOs are 12mer, with the biophysical measurements favoring a triple-layer four-fold (3×4) model and double-layer 6-fold (2×6) model over a single-layer 12-fold ring.^23^ In order to test our hypothesis we have therefore compared in another round of molecular dynamics simulations the stability of these two geometries, either bound with lauric acid or without. Consistent with the experiments of the Rangachari Lab, we chose again a ratio of 1:1 for lauric acid molecules to Aβ_42_-chains, i.e., twelve lauric acid molecules are added to the solvated fibril models. The results of these simulations are shown in Fig. 7. Interestingly, we found in all four trajectories that the (3×4) models (Fig. 7 (A)) bound with lauric acid are more stable than even the best cases seen in the control run (black line) of the model without lauric acid, but the strengthening effect is less than what we found in our previous runs for the triple-layer two-fold model. This difference is not because of any strains introduced by the ring geometry as we see in Fig. 7 (B) an even smaller stabilization for the (2×6) ring model. This is surprising as our previous work^23^ and atomic force measurements of pore size, vertical and horizontal dimensions appear to favor a double-layer ring made of six chains each rather than a triple-layer ring made of four chains.

**Figure 7.**
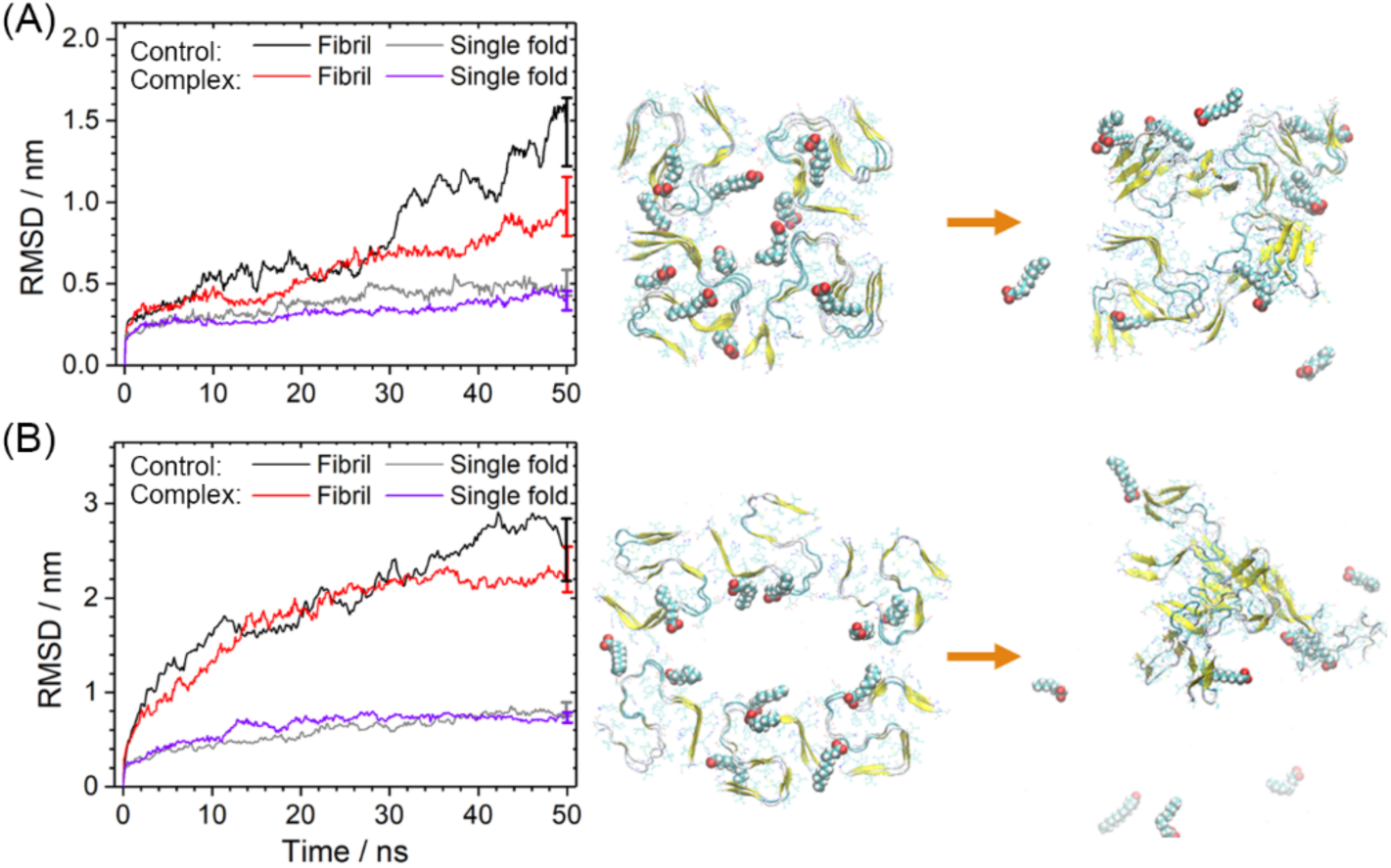
RMSD with respect to the start configuration as function of time. Date are averages over four independent trajectories. Drawn are data for the full system and such considering only single folds. Shown are also representative initial and final configurations. Results for the (3×4) model are presented in (A) and for the (2×6) model in (B).

## Conclusions

Using molecular dynamics simulations, we have studied the effect of fatty acids such as lauric acid on the stability of fibril fragments built from either U-shaped Aβ_40_-peptides or from triple-stranded S-shaped Aβ_42_-peptides. Our first result is that the addition of lauric acid does not strengthen the assembly of the sole patient derived fibril model deposited in the Protein Data Bank. This suggests that the observed lack in fibril polymorphism in patient brains is not a consequence of a stabilization of these geometries by fatty acids in the brain environment. On the other hand, a number of recent experiments have shown that fatty acids such as lauric acid can catalyze formation of Aβ assemblies.^18^ An example are the Large Fatty Acid-derived Oligomers (LFAOs) that have been studied extensively by the Rangachari Lab.^27,28^ These oligomers are built from Aβ_42_-peptides which unlike Aβ_40_-peptides do not lead to fibrils built from U-shaped chains but rather yield fibrils made of three-stranded S-shaped chains. Given the differences in chain geometries, we have therefore also studied the effect of lauric acid on the stability of various Aβ_42_ - assemblies. For these aggregates we observe indeed a noticeable strengthening by adding of lauric acid. Our results seem to indicate that lauric acid encourages elongation of the single-fold protofibrils rather than arrangement of the triple-stranded S-shaped chains into multi-fold assemblies. Especially, we see only a small stabilization for the ring-like assemblies proposed by us for the LFAOs studied by the Rangachari Lab. Hence, we conjecture that lauric acid rather plays a role in elongation of Aβ_42_ – assemblies than in their nucleation.

## Author Contributions

All four authors contributed to data generation, data analysis, and writing of the manuscript.

## Supplementary Material

Time evolution and representative initial and finial conformations of the control runs for the four Aβ_40_ fibril models (Figure S1). Root-mean-square deviation (RMSD) and representative conformation of single and double-layer two-fold Aβ_42_ models (Figure S2).

## Acknowledgment

We thank Dr. Erik J. Alred for the works on the early stage. Simulations were done on the SCHOONER cluster of the University of Oklahoma, the Extreme Science and Engineering Discovery Environment (XSEDE) under grant MCB160005, and the Blue Waters sustained-petascale computing project, which is supported by the National Science Foundation (awards OCI-0725070 and ACI-1238993) and the state of Illinois. We acknowledge financial support from National Institutes of Health (NIH) under research grants GM120634 and AG062292. E.K.V. thanks Shodor Education Foundation for all their help, and for their continued support of the Blue Waters Student Internship Program.

