## Supplemental figures for "Stability of Aβ-fibril fragments in the presence of fatty acids"

**Corresponding Author**

* Ulrich H. E. Hansmann

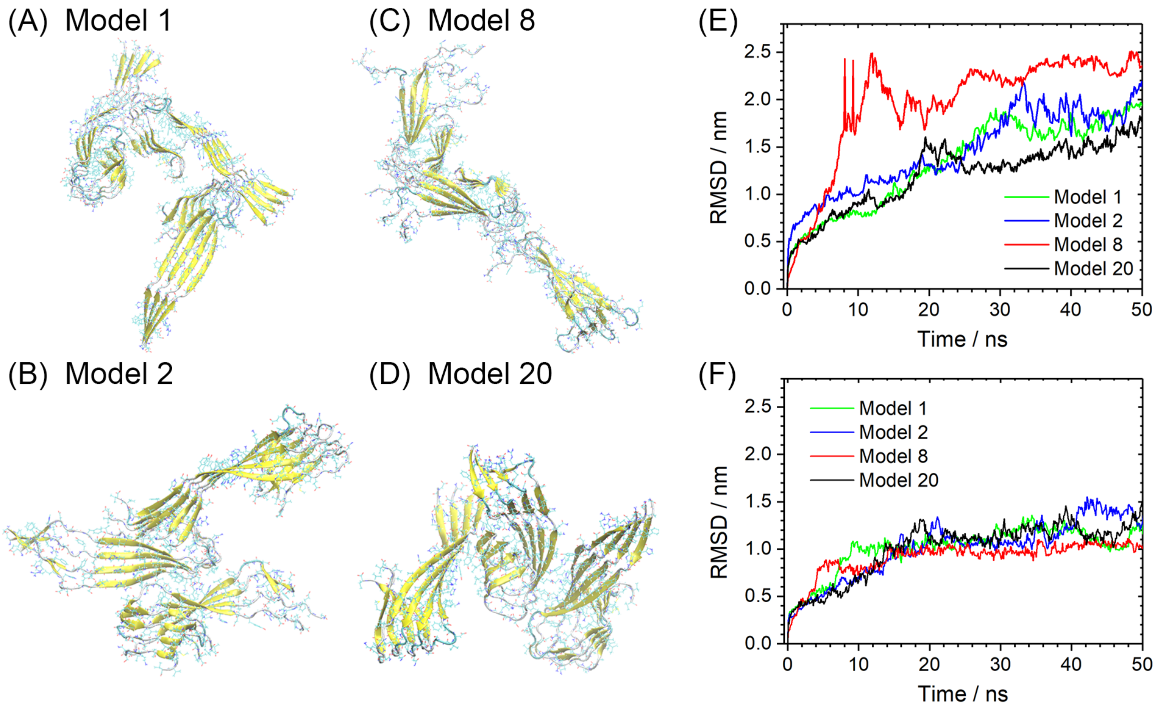


**Figure S1.** Representative final conformations of the control runs for the four models are shown in sub-figures (A-D). Root-mean-square deviations (RMSD) to the start configurations as function of time are shown in (E) where the RMSD is calculated for the whole fibril fragment. The shown values are averages over four independent trajectories. A similar plot is shown in (F) where the RMSD is calculated only over single chains (and averaged over all chains).

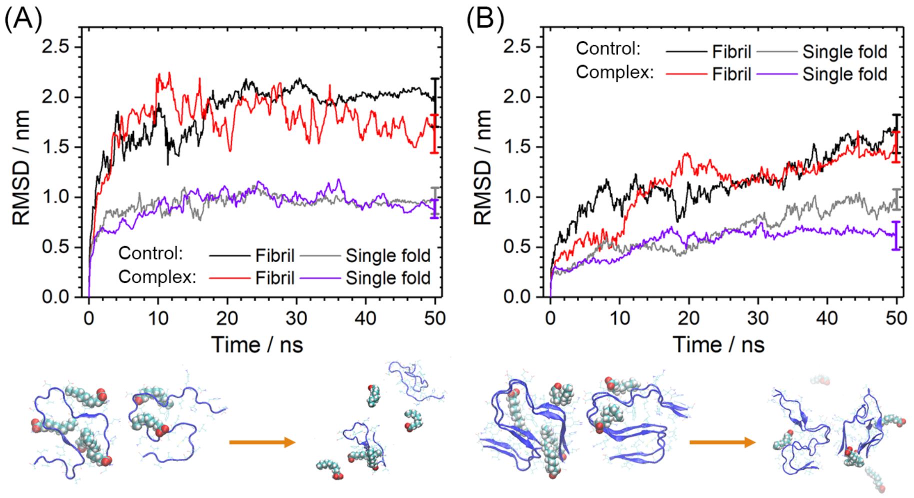


**Figure S2**. Root-mean-square deviation (RMSD) to the start configurations as function of time. Data are averaged over four independent trajectories, and representative conformations are shown to visiualize the time evolution of the systems. Data for the single two-fold A$\beta_{42}$ model are shown in sub-figure (A), and the corresponding data for the double-layer model in (B).
